# Highly selective Cooperative Primers for multiplex detection of rare mutations

**DOI:** 10.1101/2021.07.23.453558

**Authors:** Jana Oliveriusova Kent, Masen Chad Christensen

## Abstract

We describe a real time PCR-based technique capable of detecting and quantifying rare somatic mutations in circulating tumor DNA reference materials. Our approach utilizes previously described Cooperative Primers, structurally modified to exhibit high allele-specificity and one-copy target sensitivity. Cooperative Primers are bi-functional molecules, consisting of a high affinity probe fragment that guarantees sensitivity, and a covalently attached lower affinity primer providing specificity. Additional optimization of Cooperative Primer structure generated molecules capable of reliable detection of allele changes as small as a single nucleotide. These highly selective Cooperative Primers maintain excellent discrimination properties in rare mutant allele scenarios, in both monoplex and multiplex assays. With synthetic DNA samples, Cooperative Primers can detect as little as 100 copies of mutant template amongst 1 000 000 copies of wild-type template (minor allele fraction of 0.01 %). Multiplex Cooperative Primer assay was validated with cell-free DNA reference materials and consistently detected the lowest minor allele fraction available (0.1 %) for *EGFR* L858R, G719S and V769-D770insASV mutations, while simultaneously providing qualitative and quantitative assessment of cell-free DNA with integrated *β-Actin* assay. Easy to design, rapid and inexpensive, Cooperative Primer - based real time PCR assays are a promising tool for evaluation of cancer therapy response, occurrence of resistance mutations and relapse monitoring.

## Introduction

In the past few years, considerable efforts have been directed toward the development and validation of methods for gene mutation profiling of liquid biopsy samples [1-13]. Liquid biopsy, a simple and non-invasive alternative to surgical biopsies, is a source of a variety of biomarkers including circulating tumor cells, circulating tumor DNA (ctDNA), circulating miRNAs, and tumor-derived extracellular vesicles [14-17]. Circulating tumor DNA originates from apoptotic and necrotic tumor cells that shed their DNA content into the circulatory system [3, 18-21]. The analysis of ctDNA from blood offers a range of clinical application such as detecting cancer in early stages, monitoring tumor burden in response to treatment, identifying occurrence of resistance mutations and detecting disease relapse [22].

As promising as the ctDNA analysis in liquid biopsy is, it faces numerous challenges. Since normal apoptotic cells shed their DNA into the bloodstream as well, the ctDNA originating from tumor cells is very rare, often representing as little as 0.01% of the total cfDNA. Additionally, the cell-free DNA (cfDNA) is extensively fragmented, averaging about 160 bp in length [23, 24] and exhibiting a modest half-life ranging from 16 minutes to 2.5 hours [25].

Several techniques capable of overcoming these challenges have been developed to detect and analyze ctDNA [26]. Next generation sequencing (NGS) is a very powerful large-scale DNA sequencing technology allowing researchers to sequence and analyze data at a rate previously not possible. It can identify a variety of cancer-associated alterations such as point mutations, (genome wide) copy number variations, small insertions and deletions, large genome rearrangements and methylation changes, thus making it an excellent approach for cancer screening and early diagnosis [27, 28]. However, NGS library preparation, sequencing and data analysis are labor-intensive processes requiring experience. Furthermore, high cost and high random error rate render NGS unsuitable for reliable detection of minor allele fraction (MAF) < 0.1 % [29]. The poor sensitivity can be mitigated by employing a deep sequencing approach, which further increases the cost [30].

Various PCR-based techniques for detection of minority alleles have been developed and have been reviewed in detail elsewhere [22, 26]. Although typically limited to interrogation of one, or a small number of known somatic alterations, some PCR-based methods offer excellent sensitivity and fast turnaround time. The low cost and ease of use provided by real-time PCR methods make them great tools for monitoring cancer recurrence and treatment efficacy. Real-time PCR assays for identification of somatic mutations can be divided into three groups. The first group consists of techniques relying on primers that exhibit high allele-specificity, enabling them to selectively amplify mutant allele, while largely ignoring the wild-type template, such as Amplification Refractory Mutation System (ARMS) [31, 32], SNPase-ARMS [33], myT PCR [34], DPO PCR [35] and Superselective primers [36]. The second group employs various wild-type-blocking techniques to suppress the amplification of the wild-type template, while permitting amplification of the rare mutant allele such as WTB-PCR [37], PNA-LNA PCR Clamp [37], AS-NEPB PCR [38] and Snapback primers [39, 40]. The last group of assays employs modified PCR conditions, for example temperature cycling, resulting in preferential amplification of mutant allele, such as COLD-PCR[41].

Two primer designs included in the abovementioned groups structurally resemble Cooperative Primers (CoPrimers). Superselective primers are built with two oligonucleotides, mutually connected by a “bridging” sequence. The relatively long 5’ sequence segment (termed “anchor”) hybridizes to target DNA fragments with high affinity, while the shorter, 3’ segment (termed “foot”) is sensitive to the presence of variants within its binding region. The anchor and foot segments are connected by a bridging oligonucleotide, consisting of a nucleic acid sequence that has no complementarity to the foot-anchor intervening region. The foot is designed to be a 100 % match to the mutant target but mismatches the wild-type sequence; with the variant-induced mismatch positioned at its ultimate (3’ end) of the or penultimate (second base from the 3’ end) nucleotide. The bridge, lacking complementarity to the target sequence, forms a single-stranded bubble, thus effectively separating the primer’s polymerization function from the target hybridization function. As the foot is very short, it is rather inefficient in binding to the template, which is further potentiated by the presence of the (pen)ultimate base mismatch. As a result, Superselective primers almost exclusively amplify the matched (mutant) templates, while largely failing to amplify the mismatched (wild-type) template [36]. The selectivity of this system can be further enhanced by inclusion of tetramethyl ammonium chloride or bis-tetramethyl ammonium oxalate in the PCR reaction [42]. With these specificity-enhancing additives, Superselective primers are able to detect rare point mutations down to MAF of 0.025 % while using linearized wild-type and mutant plasmid template mixtures, extracts of human plasma spiked with linearized mutant plasmid, or mixtures of digested human genomic DNA extracted from normal and homozygous mutant cell lines [42].

The Snapback PCR primers also consist of two fragments - a regular PCR primer, and a probe element[39]. The probe, an oligonucleotide extension of the PCR primer, is complementary to a region of choice within the resulting amplicon. During the PCR, the probe element hybridizes to the extension product, while forming a hairpin. The stability of the hairpin is dependent on the length of the hairpin loop (distance of the primer from the probe binding region) and on the melting temperature of the stem (complementary part of the probe element). If designed appropriately, the hairpin formation hinders the Snapback primer’s extension, thus resulting in allele bias. This bias, further augmented by fast cycling conditions, has successfully been used for rare allele detection down to 0.1 % in case of point mutations, or 0.02 % for larger deletions [40].

We investigate the use and performance of CoPrimer technology [43, 44] for rare allele detection and quantitation. CoPrimers consist of a short priming sequence and a more stable capture sequence. Priming and capture fragments are connected by a non-extendable flexible linker (Fig 1). The capture sequence, due to its higher affinity, hybridizes to the template first. As a consequence, the priming sequence is brought into vicinity of its target sequence, thus increasing the local concentration of the primer by a factor of ~1 500 [44]. Finally, the priming sequence hybridizes to the template, allowing extension from its’ 3’ end. As the priming sequence is designed to bind upstream of the capture, the latter can double (upon labeling with a fluorophore and a quencher) as a hydrolysis probe. Because the capture sequence must first bind for the short primer to extend, theoretically 100 % of primer extensions result in probe cleavage, leading to high signal-to-noise ratio [43].

**Fig 1.**
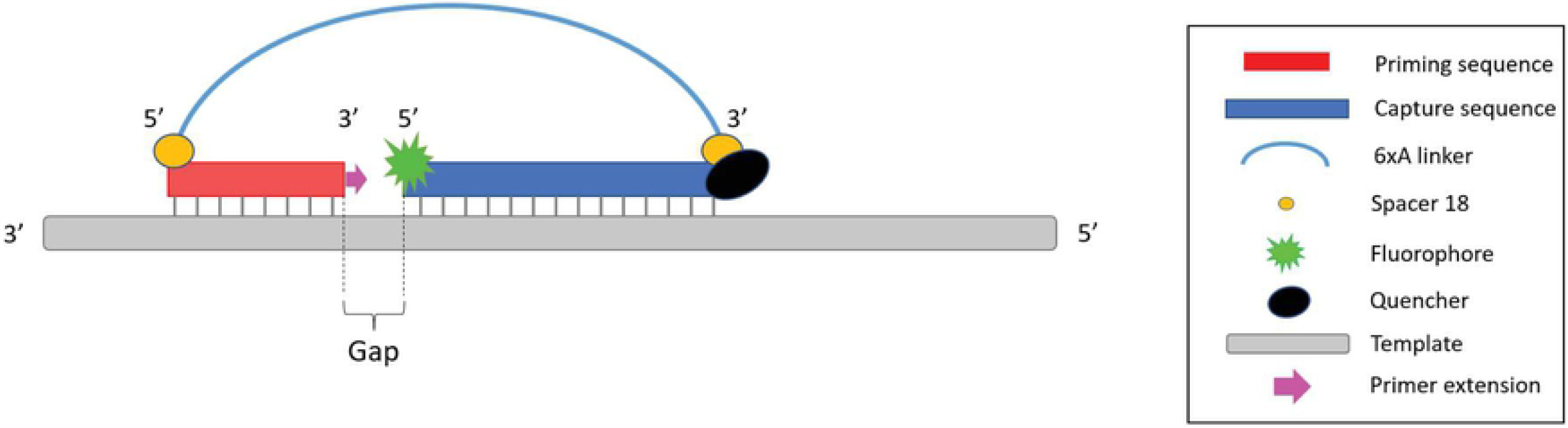
Schematic representation of CoPrimer design. CoPrimer molecule comprises of two fragments, one with higher affinity for the template (capture sequence), and one with lower affinity (priming sequence). The two building blocks of CoPrimer are joined by a flexible linker. The linker is blocked on both ends by Spacer 18 molecules to prevent DNA polymerase from extending through the linker. The capture sequence is labeled with a fluorophore and a quencher. The priming sequence functions as a primer and can be extended by DNA polymerase from its 3’ end. During primer extension, the capture sequence is hydrolyzed, and the fluorophore is released.

The priming sequence exhibits robust performance in the PCR despite its short length, because the hig-affinity capture sequence enhances primer binding stability. However, random binding of the priming sequence in the absence of pre-requisite capture hybridization is highly unlikely. Low priming fragment affinity combined with polymerase propagation inhibition due to the presence of the non-extendable linker renders CoPrimers largely immune to primer-dimer formation and propagation [44]. As such, CoPrimers are not only amenable to multiplexing, but also eradicate false-positive results due to primer-dimer formation.

Owing to its abbreviated length, the primer sequence provides an opportunity for design of highly selective CoPrimers. Early work on allele-specific CoPrimers [43] investigated the effect of variant-induced mismatch location and priming and capture Tm on the CoPrimer differentiation power. Several conclusions were made from these experiments. First, the allele-differentiation power (expressed as difference in cycle treshold, or dCq) of CoPrimers with the variant-induced mismatch positioned at the ultimate base of the priming sequence was approximately 7.1 cycles. This value could be somewhat increased (dCq of 7.6) by placing an intentional (additional) mismatch at the antepenultimate base of the priming sequence. The placement of the variant-induced mismatch under the capture sequence resulted in only modest allele differentiation power (dCq 1.7-5.3). Second, the CoPrimer designs exhibiting the greatest allele-specificity possessed priming and capture sequences of Tms of 5 °C to 7 °C, or 4 °C to 6 °C below the reaction temperature, depending on the linker length [43]. In the current publication, we explored additional ways to modify the CoPrimer structure in order to enhance its differentiation power. Our findings demonstrate the utility of CoPrimers in real time PCR assays for rare allele detection. CoPrimers exhibit not only excellent allele differentiation properties, but also possess great target sensitivity, while their resistance to primer dimer formation further decreases the likelihood of false positive result. Their amenability to multiplexing in rare allele scenarios is an added bonus, as multiplexed assays are less labour intensive and more cost effective.

## Materials and Methods

### CoPrimers

CoPrimers used in the experiments were designed with the proprietary CoDx Design Software (Co-Diagnostics Inc., Salt Lake City, UT) and synthesized by LGC Biosearch Technologies (Petaluma, CA). All CoPrimers were reconstituted in 10mM TrisHCl, 1mM EDTA, pH 8.0. The CoPrimers responsible for allele differentiation (forward or reverse) were labeled with a fluorophore and a quencher; the non-differentiating CoPrimer in each primer pair remained unlabeled. All CoPrimers used in this study are listed in Tables 1 and 2.

**TABLE 1.**
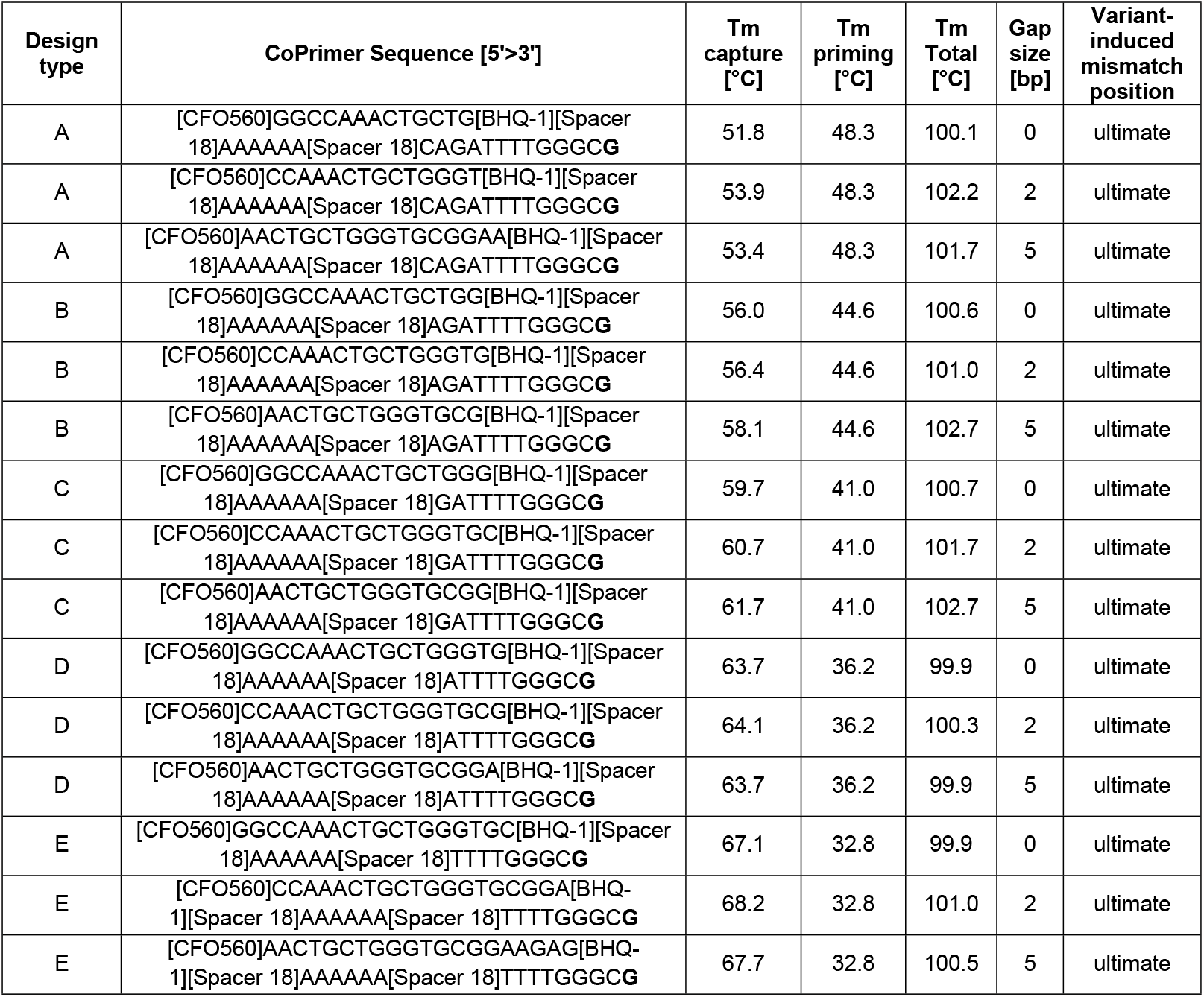
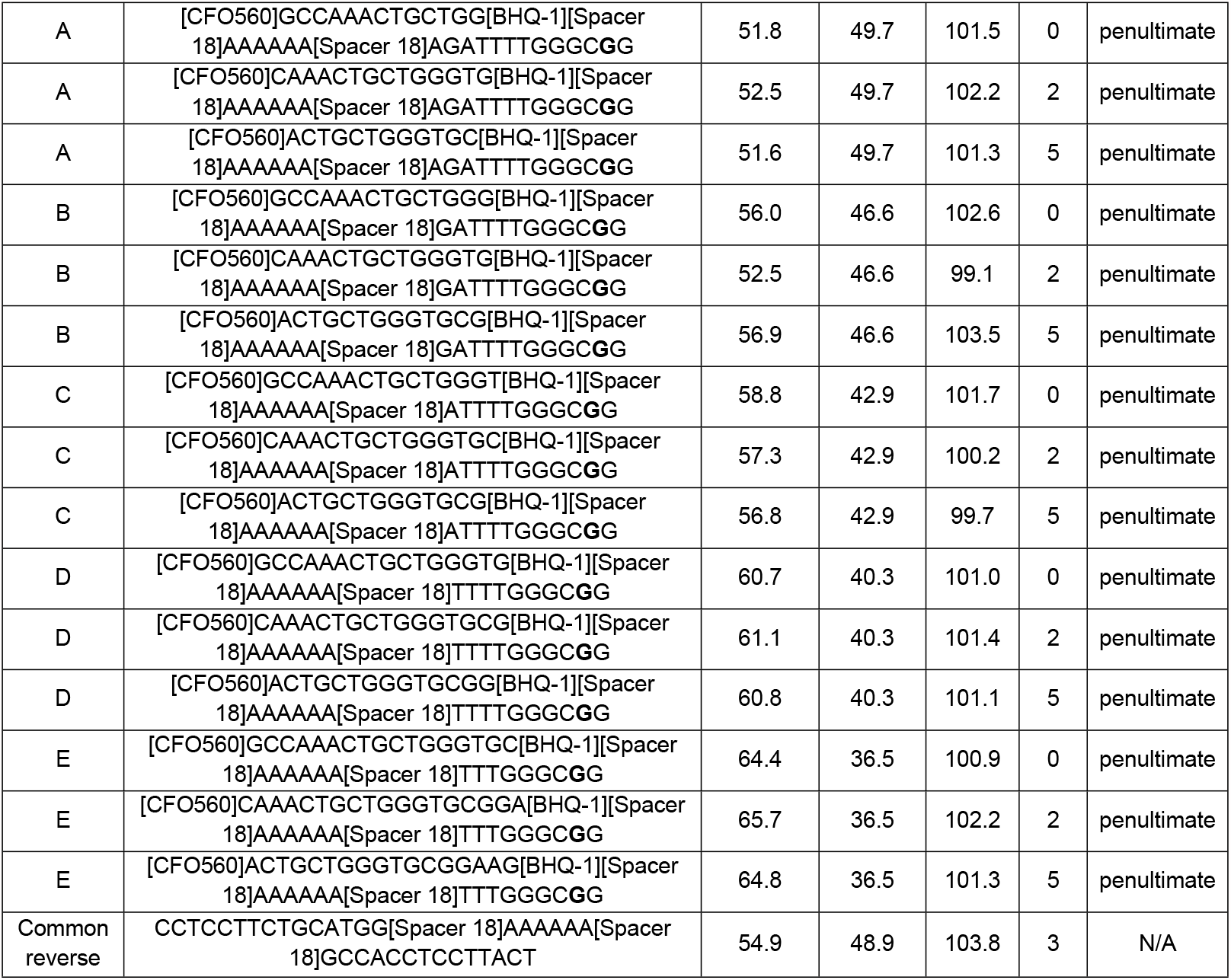
List of *EGFR* L858R-specific CoPrimers used in the design parameters optimization study. The variant of interest (L858R; c.2573T>G) is shown in bold.

**TABLE 2.**
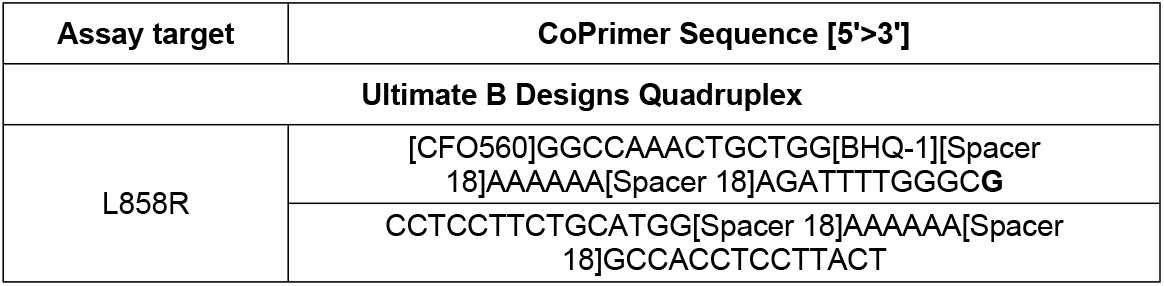
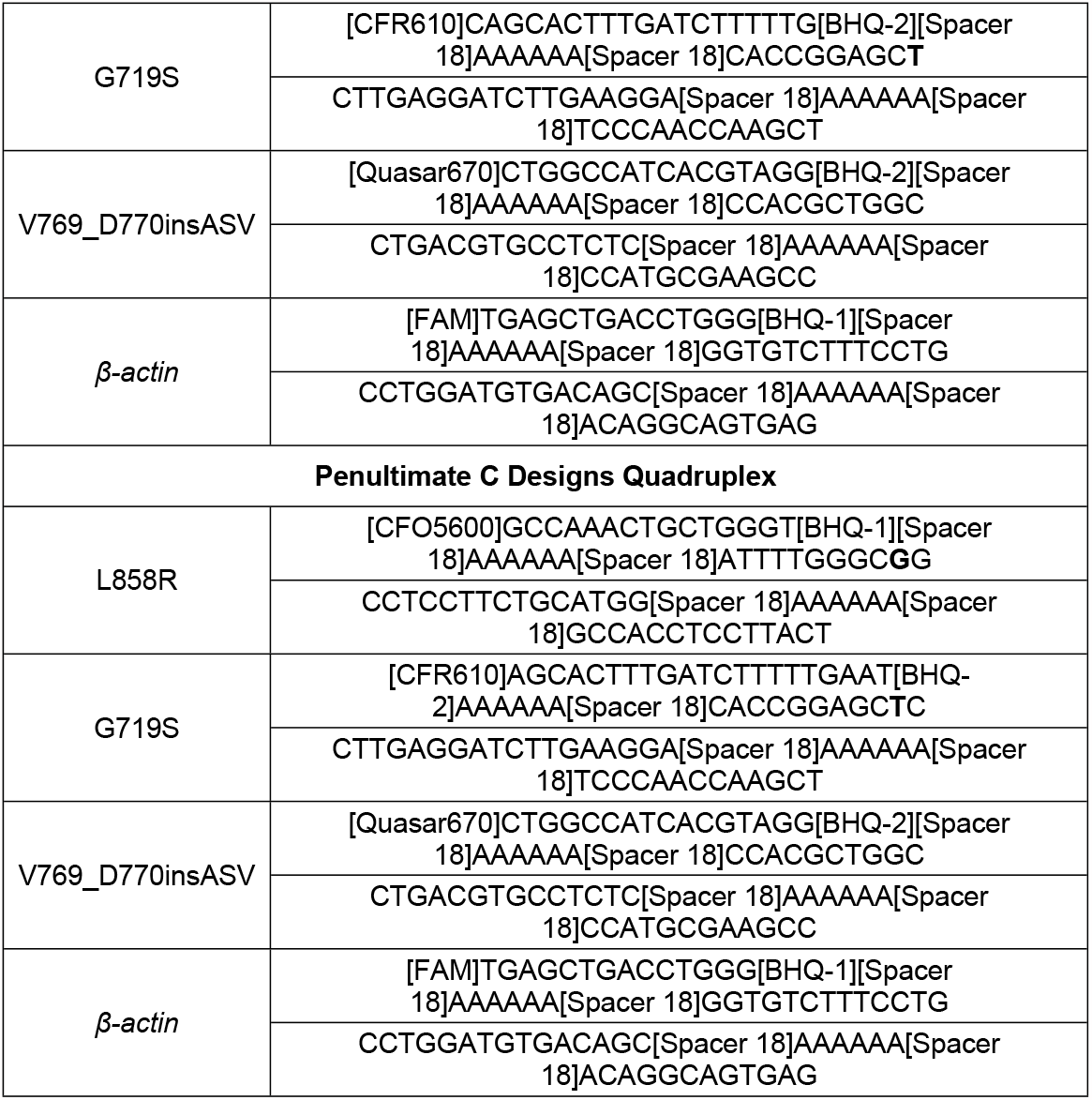
List of CoPrimers used in the quadruplex assays. The variant of interest (L858R; c.2573T>G) is shown in bold.

### Synthetic DNA templates

Synthetic double stranded human *β-actin* DNA template and *EGFR* wild-type and mutant DNA gBlock templates were ordered from Integrated DNA Technologies (Coralville, IA). To simulate the fragmented nature of cfDNA, these gBlock templates were designed to be 160bp in length. All synthetic templates were reconstituted in 10mM TrisHCl, 1mM EDTA, pH 8.0.

### Reference materials

The *EGFR* Multiplex cfDNA Reference Standard Set was purchased from Horizon Discovery (Cambridge, UK). This set is supplied at 5 %, 1 %, 0.1 % and 0 % minor allele frequencies and covers ten *EGFR* variants associated with responsiveness to *EGFR* tyrosine kinase inhibitors. While derived from normal and cancer cell lines and fragmented to an average size of ∼160 bp, it closely resembles cfDNA extracted from human plasma.

### PCR conditions

All assays, monoplexed or multiplexed, consisted of 1x ROX-free BHQ Probe Master Mix (LGC Biosearch Technologies, Teddington, UK), 0.1 μM mutant-specific, fluorescently labeled CoPrimer and 0.1 μM unlabeled common primer (Tables 1 and 2). PCR was performed in 10 μL reaction volumes consisting of 7 μL of the master mix combined with 3 μL of either the synthetic DNA template or the reference material. Each PCR reaction was mixed by pipetting up and down prior to temperature cycling, to ensure homogeneity of assay contents. The thermocycling was performed in 4-channel MIC real-time PCR instrument (Biomolecular Systems, Upper Coomera, Australia) using MIC 4-tube strips. The cycling protocol consisted of initial denaturation step (activation of chemically-blocked polymerase) for 15 min at 95 °C, followed by 50 cycles of denaturation step at 95 °C for 15 s and annealing/extension step at 55 °C for 60 s. The fluorescence intensity of the dyes released from the capture sequence was recorded at the conclusion of each annealing/extension step.

## Results

### Optimization of allele-specific CoPrimer capture and priming sequence length, gap size and location of the variant-induced mismatch

Each CoPrimer molecule (Fig 1) possesses several attributes: (I) affinity of the capture and priming sequences to the template, expressed as respective Tm values; (II) length and composition of the inert linker, connecting the priming and capture fragments and (III) length of the gap, characterized as the distance created between the capture and priming fragments upon their binding to the target sequence. Allele-specific CoPrimers have one additional characteristic - the location of the variant-induced mismatch within the CoPrimer molecule.

Since the publication that introduced CoPrimers in 2014 [43], a modification has been made to CoPrimer structure with regards to the linker composition. As the original linker typically consisted of 4-6 Spacer 18 molecules, the CoPrimer synthesis was a rather costly process. It has been hypothesized and subsequently proven that a linker consisting of two Spacer 18 molecules with an intervening polyA sequence of six nucleotides significantly decreases the synthesis cost, and maintains both PCR efficiency and primer-dimer resistance when compared to the original design (unpublished results). To define the optimal lengths of priming and capture sequences associated with the highest allele discrimination properties, we designed a series of forward CoPrimers with varying lengths of capture and priming fragments (Table 1), while keeping the linker at a constant length of two Spacer 18 molecules with six intervening adenosine residues. These designs were designated as A, B, C, D and E. While the A designs have the longest priming sequence and the shortest capture; the E designs possess the longest capture and the shortest priming sequence (Table 1).

To evaluate the impact of the interrogating nucleotide location on allele-specificity, CoPrimers with varying priming and capture fragment lengths were designed in two iterations: with variant-induced mismatch located either at the ultimate base, or at the penultimate base of the priming sequence. For CoPrimers with the mutation-induced mismatch at their ultimate base, the priming sequence was designed to vary in Tm from 6.7 °C to 22.2 °C below the annealing/extension temperature; the capture segment Tm varied from 3.2 °C below to 13.2 °C above the annealing/extension temperature (Table 1). For CoPrimers with the mutation-induced mismatch at their penultimate base, the priming sequence was designed to vary in Tm from 5.3 °C to 18.5 °C below the annealing/extension temperature, while the capture segment Tm varied from 3.4 °C below to 9.8 °C above the annealing/extension temperature (Table 1). To ensure near-constant affinity of all CoPrimer designs for the template, the sum of the priming and capture Tm was kept constant as much as possible (Table 1). To investigate the influence of gap size on CoPrimer performance, each of the CoPrimer designs was synthesized with 0 bp gap, 2 bp gap and 5 bp gap.

One common reverse CoPrimer was designed to be paired with all of the different forward designs, creating amplicons varying in size between 76 and 80 bp, depending on the length of the forward CoPrimer priming sequence and the length of the gap.

The CoPrimers were designed to selectively amplify the L858R (c.2573T>G, COSM6224) mutation in *EGFR* gene. The exon 21 L858R point mutation, together with the numerous exon 19-deletions, represent nearly 85 % of all *EGFR* aberrations in patients with non-small cell lung cancer [45-47]. The *EGFR* mutation status is a valuable predictive factor regarding the effectiveness of tyrosine kinase inhibitiors that target this aberrant oncogene [48].

Allele-specificity of these CoPrimer designs was first evaluated in monoplex PCR reactions containing either wild-type or mutant synthetic template at a concentration of 1 000 000 copies/μL. The differentiation power of individual CoPrimers was expressed as dCq (Cq value obtained with mutant template *minus* the Cq value obtained with wild-type template) and listed in Table 3. We observed clear dependence of dCq values on the length of the priming sequence - the shorter the priming sequence, the larger was the dCq value, indicating increasing wild-type *vs*. mutant differentiation properties of the CoPrimer. Interestingly, this trend was significantly more pronounced in the CoPrimer designs with the interrogating nucleotide located at the ultimate base of the priming sequence, when compared to those with the variant positioned at the penultimate base. Penultimate base designs with medium to short priming and medium capture sequences (A-C designs) exhibited better allele differentiation than equivalent (A-C) ultimate designs. However, the penultimate designs gained only modest dCq improvements with further abbreviated priming sequence (D and E designs), while the ultimate D and E CoPrimer designs exhibited major dCq gain. Both ultimate and penultimate designs possessed the best discriminatory properties when priming and capture sequences were juxtaposed (gap length equal to 0 bp). Increasing the gap size to 2-5 bp was associated with poorer discrimination for both types of designs (ultimate and penultimate), with similar dCq values for 2 bp and 5 bp gaps. Overall, the ultimate design with the longest capture, the shortest priming sequence and 0 bp gap (design type D) exhibited the highest dCq values of all CoPrimers (19.0 cycles).

**TABLE 3.** Allele discrimination properties of *EGFR* L858R-specific CoPrimers.

| Design type | Ultimate base mismatch |  |  | Penultimate base mismatch |  |  |
| --- | --- | --- | --- | --- | --- | --- |
|  | Tm priming [°C] | Gap size [bp] | dCq | Tm priming [°C] | Gap size [bp] | dCq |
| A | 48.3 | 0 | 12.4 | 49.7 | 0 | 15.3 |
| B | 44.6 | 0 | 16.3 | 46.6 | 0 | 14.8 |
| C | 41.0 | 0 | 16.0 | 42.9 | 0 | 16.5 |
| D | 36.2 | 0 | 16.5 | 40.3 | 0 | 16.1 |
| E | 32.8 | 0 | 19.0 | 36.5 | 0 | 16.5 |
| A | 48.3 | 2 | 8.2 | 49.7 | 2 | 10.2 |
| B | 44.6 | 2 | 9.4 | 46.6 | 2 | 10.5 |
| C | 41.0 | 2 | 10.3 | 42.9 | 2 | 11.7 |
| D | 36.2 | 2 | 12.4 | 40.3 | 2 | 13.2 |
| E | 32.8 | 2 | 15.1 | 36.5 | 2 | 14.1 |
| A | 48.3 | 5 | 9.0 | 49.7 | 5 | 10.4 |
| B | 44.6 | 5 | 9.3 | 46.6 | 5 | 10.6 |
| C | 41.0 | 5 | 11.0 | 42.9 | 5 | 12.7 |
| D | 36.2 | 5 | 12.2 | 40.3 | 5 | 13.5 |
| E | 32.8 | 5 | 16.1 | 36.5 | 5 | 14.9 |

Next, we tested the ability of all CoPrimer designs to detect L858R mutant targets in abundant wild-type background in simulated monoplex rare allele PCR reactions while using synthetic template mixtures of known mutant allele fraction. Whilst keeping the wild-type template copy number at a constant value of 1 000 000 copies/μL in each sample, the mutant template copy number was either 0 copies/μL, 10 copies/μL, 100 copies/μL, 1 000 copies/μL, 10 000 copies/μL, 100 000 copies/μL or 1 000 000 copies/μL. A negative PCR reaction containing no *EGFR* template was included as well. All reactions were performed in duplicate. The goal of this experiment was to identify the optimal CoPrimer design possessing high sensitivity, maximum selectivity and also exhibiting satisfactory linearity (assessed by plotting Cq value against the logarithm of the mutant template copy number in the sample) over a wide span of MAF values. The results of this experiment are shown in Fig 2 (for the CoPrimer designs with the gap size of 0bp, which had the best performance).

**Fig 2.**
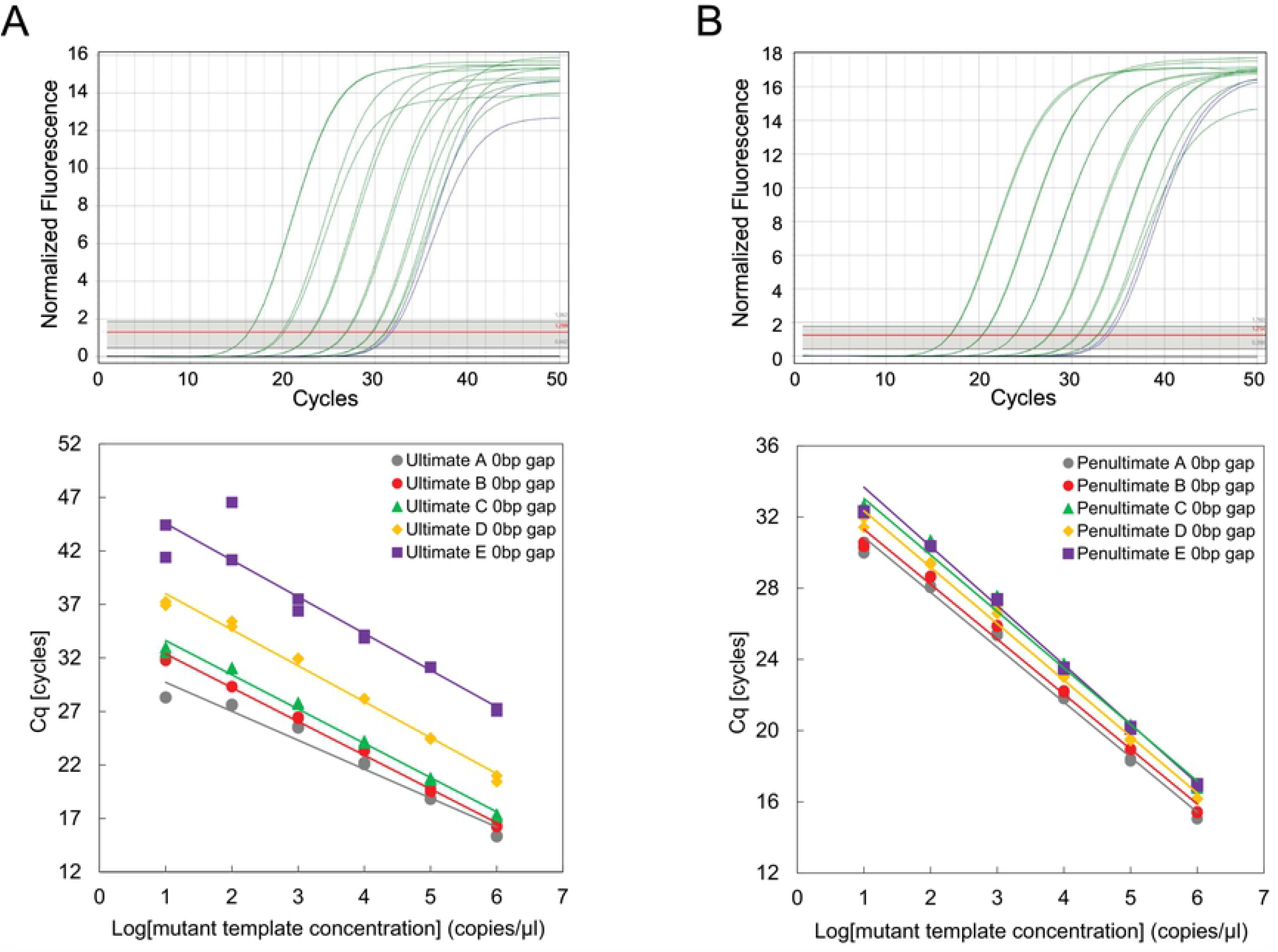
Amplification data illustrating the influence of CoPrimer priming and capture fragment lengths and location of the interrogating nucleotide on L858R CoPrimer allele-specificity in a monoplex assay. Two sets of forward CoPrimers with 0 bp gap and varying priming and capture lengths were designed to selectively amplify *EGFR* L858R mutant template. The CoPrimer designs were termed A though E, with the A designs possessing the longest priming sequence and the shortest capture and E designs having the shortest priming sequence and the longest capture. The interrogating nucleotide was positioned either on ultimate (top, A) or penultimate (top, B) base. The allele-specificity was assessed through the linearity of the Cq values on the logarithm of the mutant template concentration in the sample. While keeping the wild-type template copy number at a constant value of 1 000 000 copies/μL in each sample, the mutant template concentration was either 0 copies/ μL, 10 copies/μL, 100 copies/μL, 1 000 copies/μL, 10 000 copies/μL, 100 000 copies/μL and 1 000 000 copies/μL. Real-time amplification curves generated with ultimate B (bottom, A) and penultimate C (bottom, B) CoPrimer designs are shown. PCR reactions initiated with mixtures of 1 000 000 copies/μL wild-type template with 10 copies/μL, 100 copies/μL, 1 000 copies/μL, 10 000 copies/μL, 100 000 copies/μL and 1 000 000 copies/μL are shown in green; a wild-type-only (1 000 000 copies/μL wild-type template, 0 copies/μL mutant template) PCR reaction and negative PCR reaction containing no *EGFR* template are shown in blue and black, respectively. All reactions were performed in duplicate.

Ultimate CoPrimer designs B, C and D with 0 bp gap exhibited good linearity down to MAF of 0.001 % (R^2^ values of 0.9963, 0.9911 and 0.9906, respectively). Ultimate design A demonstrated poor linearity at MAF values lower than 0.1 %, indicating poor mutant vs. wild-type discriminatory power (Fig 2A). The ultimate E CoPrimer designs exhibited high Cq values due to poor PCR efficiency and poor Cq value reproducibility within replicates - a trend often seen with CoPrimers having priming fragments of insufficient length (unpublished observation). All CoPrimers with the interrogating nucleotide positioned at the penultimate base of the Primer and 0bp gap exhibited great linearity down to MAF of 0.001 % (Fig 2B), with the R^2^ values of 0.9914 (A), 0.9899 (B), 0.9938 (C), 0.9930 (D) and 0.9925 (E). The penultimate A-E designs maintained good PCR efficiency and consistent Cq values that were only marginally affected by the variable priming and capture fragment sizes (Fig 2B).

The influence of gap size on CoPrimer selectivity was investigated as well. The best performing CoPrimers from both ultimate and penultimate design groups were those lacking the gap between the priming and capture segments (gap size of 0 bp). Extending the length of the gap from 0 bp to 2 bp or 5 bp significantly reduced CoPrimer selectivity and negatively impacted the linearity expressed as the dependence of the PCR Cq value and logarithm of the mutant allele concentration in the sample (Fig 3). Again, the adverse effect of the gap size on selectivity was less pronounced in penultimate CoPrimer designs (Fig 3B) when compared to ultimate designs (Fig 3A).

**Fig 3.**
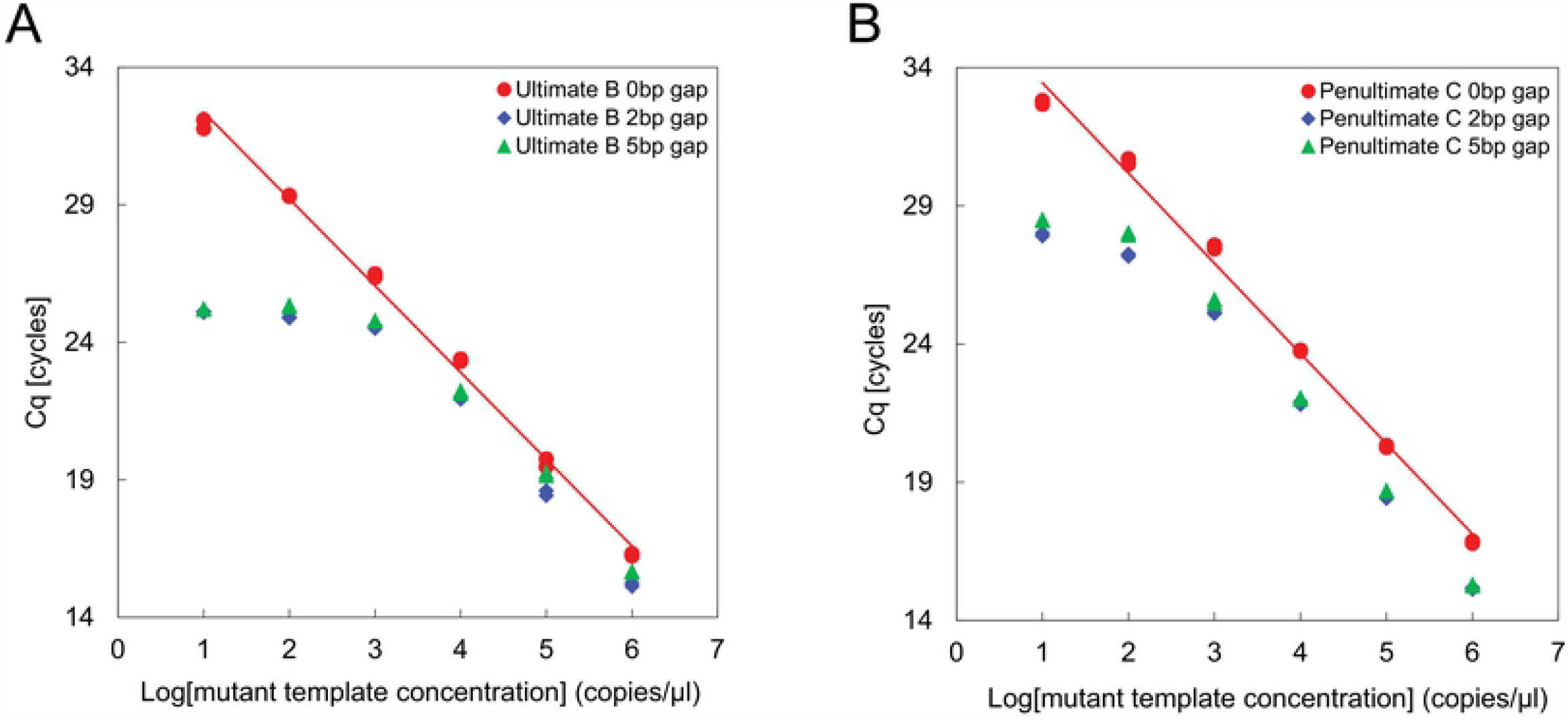
Gap length impact on CoPrimer specificity. The effect of the gap length on allele-specificity of L858R CoPrimer design B with the interrogating nucleotide at the ultimate base of the priming sequence (A), and L858R CoPrimer design C with the interrogating nucleotide at the penultimate base of the priming sequence (B) was studied using simulated rare allele samples. While keeping the wild-type template copy number in each sample at a constant value (1 000 000 copies/μL), the mutant template copy number was varied (0 copies/μL, 10 copies/μL, 100 copies/μL, 1 000 copies/μL, 10 000 copies/μL, 100 000 copies/μL and 1 000 000 copies/μL). A PCR reaction containing no *EGFR* template was included in each run as a negative control. All reactions were performed in duplicate and the Cq values for all designs were plotted against logarithm of the mutant template concentration in the sample.

### Multiplexing capabilities of allele-specific CoPrimers with synthetic DNA targets

Both the ultimate and penultimate allele-specific CoPrimer designs were also evaluated in multiplex rare-allele detection assays. In addition to the already existing L858R-specific CoPrimer pairs, two additional primers targeting *EGFR* mutations G719S (c.2155G>A, COSM6252) and V769-D770insASV (c.2307_2308insGCCAGCGTG, COSM12376) were designed. The G719S mutant-specific CoPrimer was designed using B, C and D ultimate parameters and also C, D and E penultimate parameters. The V769-D770insASV mutation is a 9 bp insertion, and was included in our quadruplex in order to determine whether the CoPrimer-based rare allele detection assay can be applied to single-nucleotide variants as well as larger aberrations such as InDels. The priming fragment of the V769-D770insASV CoPrimer was positioned over the 9 bp insertion to selectively amplify the mutant allele, while maintaining the capture/priming Tm characteristics of ultimate B, C and D and penultimate C, D and E design criteria.

To facilitate quantification of the DNA template in each reaction, another CoPrimer pair targeting a ubiquitous gene, specifically human *β-actin*, was generated to complete both ultimate and penultimate quadruplex assays. The *β-actin*-specific primer pair was designed to amplify a 93 bp fragment spanning a majority of intron 5 (nucleotides 7602-7694, NCBI reference sequence NG_007992.1). This assay design is a modification of the NSCLC cfDNA quantification assay described by Chiapetta et al.[49]. Each of the CoPrimers in the quadruplex was labeled with a unique fluorophore (Table 3). The performance of all quadruplex assays was first tested on mixtures of synthetic DNA templates for all 4 targets (3 *EGFR* mutations and *β-actin* quantification control). While keeping the wild-type template copy number in each sample at a constant value (1 000 000 copies/μL), the mutant template copy number was varied (0 copies/μL, 10 copies/μL, 100 copies/μL, 1 000 copies/μL, 10 000 copies/μL, 100 000 copies/μL or 1 000 000 copies/μL). A PCR reaction containing no DNA template was included in each run as a negative control. All reactions were performed in duplicate. Our experiments revealed that the ultimate CoPrimer design B performed robustly in this multiplex setting (Fig 4A), while ultimate designs C and D exhibited poor PCR efficiency. Similarly, the penultimate designs D and E amplified poorly in multiplex assay, while penultimate design C exhibited strong performance (Fig 4B). Both L858R and V769-D770insASV within the ultimate B assay exhibited good linearity down to MAF of 0.001 % (R^2^ values of 0.9972 and 0.9992, respectively). The L858R and V769-D770insASV within the penultimate C assay also possesed good linearity down to MAF of 0.001 % (R^2^ values of 0.9888 and 0.9974, respectively). The G719S mutation assay proved to be linear only to MAF of 0.01 %, with R^2^ values (of the data for up to MAF of 0.01 %) of 0.9960 and 0.9915 for the ultimate B and penultimate C quadruplex assays, respectively (Fig 4).

**Fig 4.**
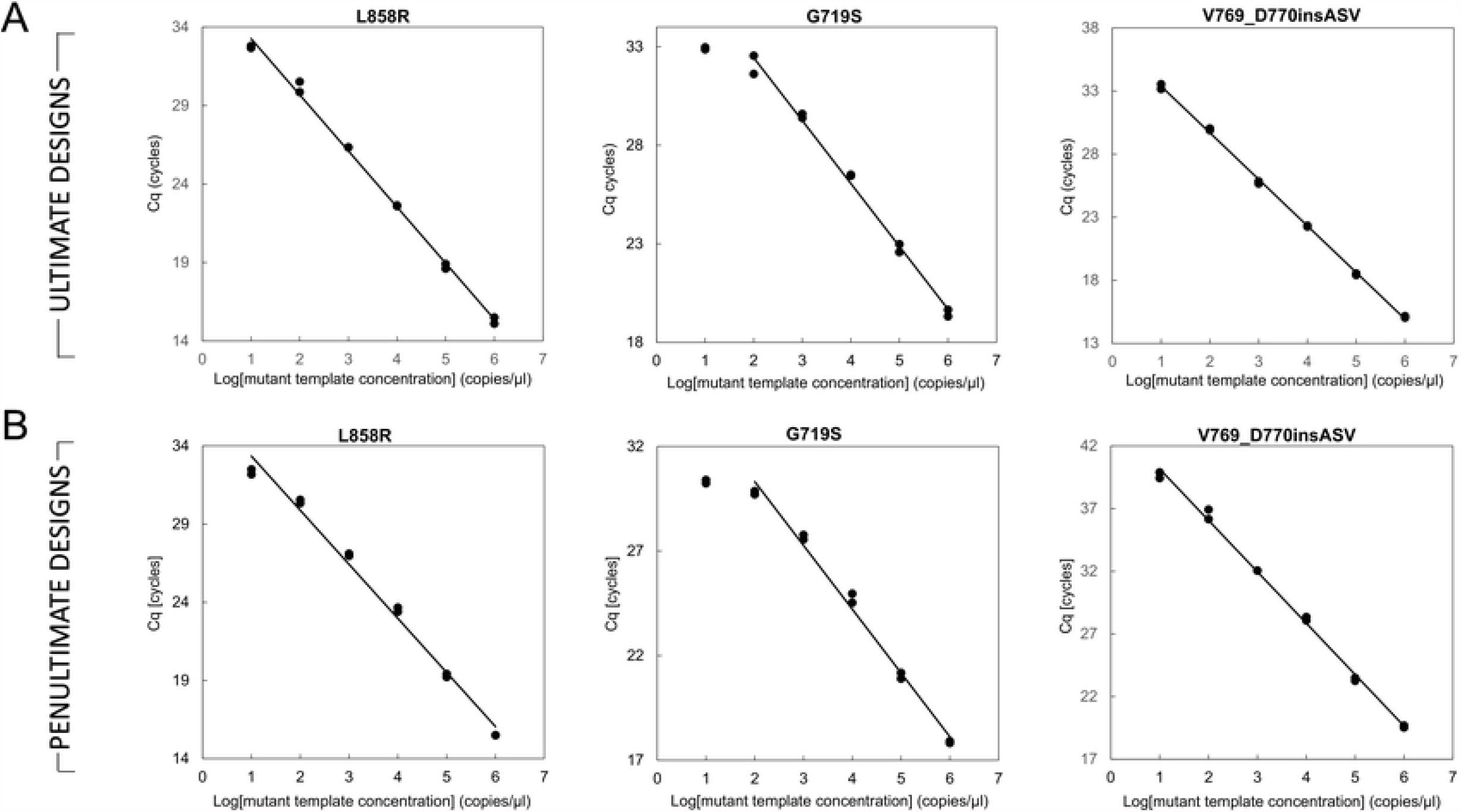
Multiplexing capabilities of allele-specific CoPrimers. Two quadruplex assays were assembled to demonstrate amenability of allele-specific CoPrimers in multiplex rare mutation detection and quantitation. Both multiplex assays contained three CoPrimer pairs detecting *EGFR* L858R, G719S and V769-D770insASV mutant templates, and a CoPrimer pair for amplification of *β-actin* quantification control. The CoPrimers constituting the quadruplex assay were either designed in accordance with the ultimate B (A, top) or penultimate C (B, bottom) design model. The PCR reactions were initiated with samples containing mixture of wild-type templates for *EGFR* L858R locus, G719S locus, V769-D770insASV locus and *β-actin* locus at a constant value of 1 000 000 copies/μL (each), while the *EGFR* L858R, G719S and V769-D770insASV mutant template copy number was either 0 copies/ μL, 10 copies/μL, 100 copies/μL, 1 000 copies/μL, 10 000 copies/μL, 100 000 copies/μL and 1 000 000 copies/μL (each). A reaction containing no template (negative control) was included as well. All reactions were performed in duplicate and the Cq values for all designs were plotted against logarithm of the mutant template concentration in the sample.

### Performance of allele-specific CoPrimers in rare-allele multiplex assays with reference materials

Using all the data points obtained from the optimization experiments, we were able to elucidate two best candidate CoPrimer designs that possess high selectivity and robust PCR efficiency in multiplex rare-allele assays: (I) the B CoPrimer design with the interrogating nucleotide positioned at the ultimate base, and (II) the C CoPrimer design with the interrogating nucleotide located at the penultimate base. The ability of these two CoPrimer designs to reliably detect rare mutant allele in abundant wild-type background was further tested using the *EGFR* Multiplex cfDNA Reference Standard Set (Horizon Discovery). Each reaction contained three CoPrimer pairs detecting *EGFR* L858R, G719S and V769-D770insASV mutant templates, in addition to the CoPrimer pair amplifying *β-actin*. Five reactions were set up for each of the two (ultimate C and penultimate B) quadruplex assays. These PCR reactions were initiated with reference samples containing the *EGFR* L858R, G719S and V769-D770insASV mutant templates at MAF of 5 %, 1 %, 0.1 % and 0 % (wild-type only). The last reaction contained no template (negative control). All reactions were performed in duplicate. The results of this experiment are shown in Fig 5, expressed as the dependence of the Cq value on the logarithm of the mutant allele concentration in the reference sample.

**Fig 5.**
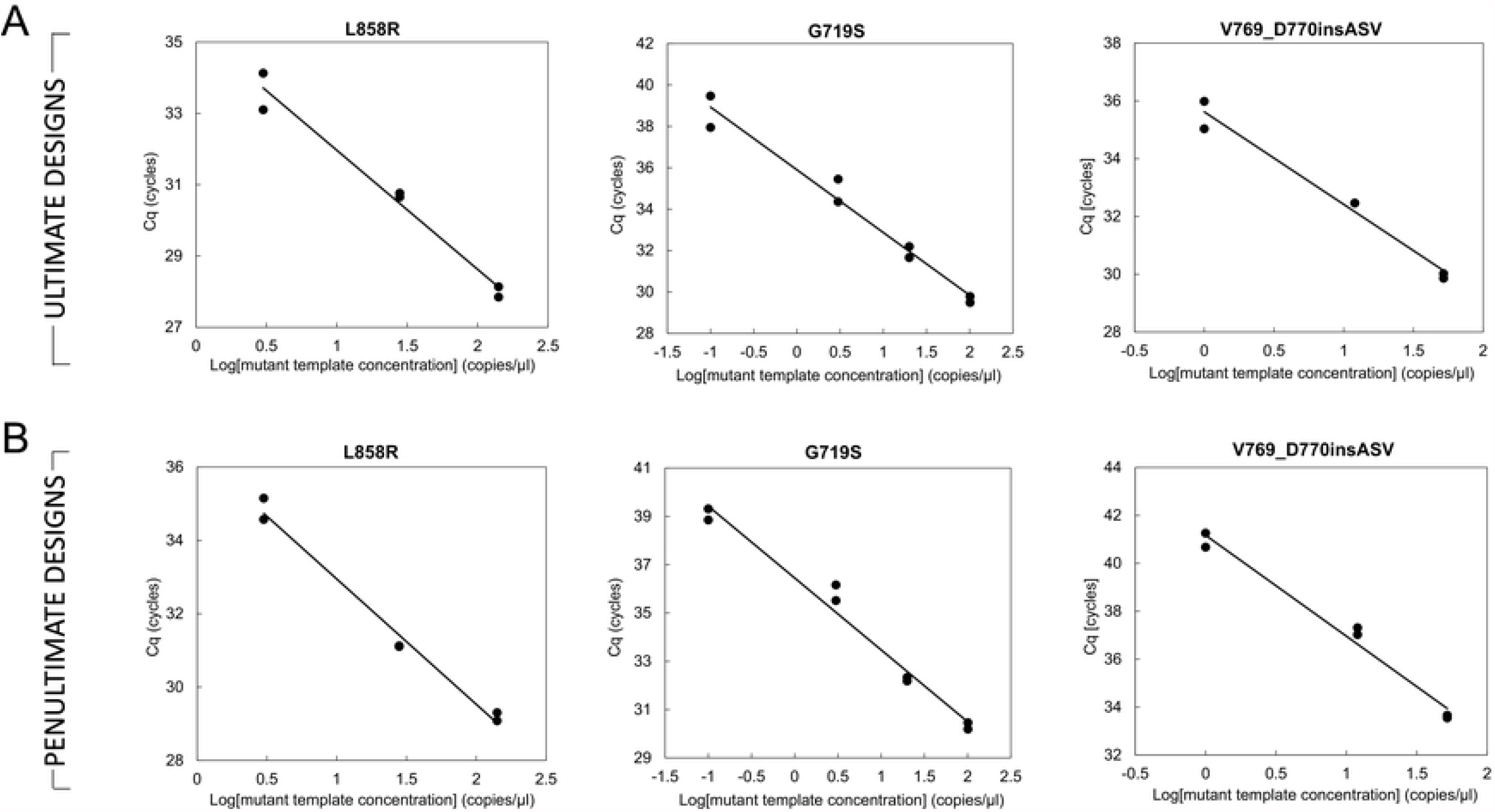
Allele-specific CoPrimers in multiplex assays on cfDNA reference materials. The performance of multiplex assays assembled from design “B” CoPrimers with the interrogating nucleotide at the ultimate base (A, top) of the priming sequence and CoPrimer design “C” with the interrogating nucleotide at the penultimate base (B, bottom) of the priming sequence were tested on *EGFR* Multiplex cfDNA Reference Standards. Each reaction contained three CoPrimer pairs detecting *EGFR* L858R, G719S and V769-D770insASV mutant templates, and a CoPrimer pair amplifying *β-actin*. The PCR reactions were initiated with a reference sample containing the *EGFR* L858R, G719S and V769-D770insASV mutant templates at MAF of 0.05; 0.01, 0.001 and 0 % (wild-type only). A reaction containing no template (negative control) was included as well. All reactions were performed in duplicate and the Cq values for all designs were plotted against logarithm of the mutant template concentration in the sample.

The ultimate C quadruplex assay showed great sensitivity and specificity for all three mutant targets (Fig 5A). Each individual assay within the multplex exhibited linear amplification response for the three reference samples with MAF varying between 5 % and 0.1 %. The wild-type-only control for the L858R mutation amplified with a significant Cq delay (15.7 cycles on average) when compared to that of the sample with the lowest concentration of the mutant DNA (MAF of 0.1 %). The wild-type-only control for the V769-D770insASV sample did not amplify at all, thus producing zero wild-type background (Fig 5A). The wild-type-only sample for G719S amplified with a significant Cq delay (9.1 cycles on average). However, according to the quality control documentation provided by the manufacturer, the G719S wild-type-only sample contained low levels of the mutant DNA (0.1 copies/μL, translating into MAF of 0.005 %). Interestingly, the Cq value for the “wild-type only” sample falls within the linear amplification response of the remaining reference samples with MAF of 5-0.1 %. It is difficult to say whether the CoPrimer amplification response for G719S “wild-type only” sample is due to the amplification of the low copy mutant target present, or whether it is a result of the non-specific amplification of the wild-type template. Very similar results were observed with the penultimate B quadruplex assay (Fig 5B). Linear amplification response was observed for all three mutant-specific (L858R, G719S and V769-D770insASV) assays over the span of MAF between 5 % and 0.01 % provided by the reference samples. The wild-type-only control for the L858R mutation amplified only in one of the two duplicates, with a Cq delay of 10.3 cycles when compared to that of the sample with the lowest concentration of the mutant DNA (MAF of 0.1 %). Similarly to the ultimate C quadruplex assay, the wild-type only reference sample for the V769-D770insASV sample did not amplify at all. The wild-type-only sample for G719S (containing low-level mutant DNA of MAF of 0.005 %) amplified with an average Cq delay of 8.8 cycles (Fig 5B).

## Discussion

Cooperative primers possess several attributes that make them uniquely suited for the development of complex real-time PCR assays. They are resistant to primer-dimer formation and produce robust, yet specific amplification. These characteristics persist in highly multiplexed assays, which have been utilized in a number of CoPrimer-based commercial molecular tests (www.codiagnostics.com).

Use of allele-specific CoPrimers for variant differentiation has been previously investigated, albeit not in depth [43]. The previous investigation used ARMS-(priming) and probe (capture) - based differentiation, but the allele discrimination power of these CoPrimer designs was not high enough for them to be suitable for use in genotyping assays. In our publication, we focused on systematic optimization of CoPrimer design parameters to maximize the molecule’s allele-specificity while maintaining the sensitivity and amenability to multiplexing. This was achieved through researching the ideal position of the variant within the priming sequence, the lengths of the priming and capture segments, and the size of the gap. This optimization was performed in two steps: with *EGFR* L858R mutation-specific CoPrimers in a monoplex assay, and with *β-actin, EGFR* L858R, G719S and V769-D770insASV-specific CoPrimers in a multiplex assay.

Non-specific hybridization of the L858R mutant-specific CoPrimer to the wild-type template results in a G:A mismatch located either at the ultimate, or penultimate base of the priming sequence, dependent on the design. The allele discrimination by CoPrimers possessing this mismatch were excellent (Table 3.). The specificity of CoPrimers was further improved by reducing the length of the priming sequence. Decreasing the priming sequence length lowers the affinity for the template (Tm), which increases the relative destabilization effect of the mismatch and as a result, increases CoPrimer specificity. Thanks to the stabilizing effect of the high-affinity capture sequence, robust PCR amplification was obtained even with CoPrimers having priming sequence Tm as low as 32.8 °C (ultimate C design), which is more than 22 °C below the annealing temperature used in the PCR. The inverse dependence of allele discrimination (expressed as dCq) on priming Tm was more apparent in ultimate designs when compared to penultimate designs (Table 3). However, the penultimate designs with longer priming fragments (designs A, B, C) exhibited greater allele discrimination power (dCq values) than the ultimate designs with the same priming sequence lengths. Consequently, the additional gain in discrimination power with further reduced priming sequence length is not as apparent in penultimate D and E designs when compared to ultimate designs. It can thus be concluded that the penultimate variant-induced mismatches in CoPrimers with longer priming fragments evoke the same duplex destabilization as ultimate mismatches in CoPrimers with shorter priming fragments. However, the greatest allele-specificity (dCq of 19.0) was achieved with the ultimate CoPrimer E design (shortest priming fragment tested), while penultimate E design exhibited dCq of only 16.5. However, this excellent allele-specificity comes at a cost. CoPrimer designs with too short of a priming sequence suffer from poor PCR efficiency, and therefore do not perform well in multiplex reactions. Poor overall performance of ultimate CoPrimers with priming sequences of a Tm ≤ 40.9 °C (ultimate designs C, D and E) and or penultimate CoPrimers with priming fragments of a Tm ≤ 40.3 °C (penultimate designs D and E) was observed in multiplex assays assembled from 3 *EGFR*-specific CoPrimer pairs and the *β-actin* CoPrimer pair. Extending the length of these (apparently too short) priming sequences by a single nucleotide increased the Tm of the priming segment to 44.6 °C and 42.9 °C (ultimate B design and penultimate C design, respectively), which fully restored CoPrimer performance in the quadruplex assay to the robust level observed in monoplex reactions.

The investigation of gap size influence on CoPrimer specificity revealed that smaller gaps conferred better CoPrimer specificity. This was not only apparent from the L858R monoplex PCR experiments (Table 3 and Figs 2 and 3), but was also evident from the performance of L858R, G719S, V769-D770insASV and *β-actin* quadruplex assays (Fig 4). The linearity of the dependence of the Cq value on the logarithm of the mutant allele concentration in the sample could be significantly improved by decreasing the CoPrimer gap size (Fig 3). The best performance was observed with CoPrimer designs with juxtaposed priming and capture fragments (0 bp gap size), and this trend was observed for CoPrimers of all priming fragment sizes (designs A, B, C, D and E) and both ultimate and penultimate designs (Fig 3). It has been theorized that gaps of lengths ≤ 3 bp negatively affect binding capability of CoPrimer to the target template, mostly due to decreased flexibility of the CoPrimer molecule [44], and limited flexibilty of the template. Lack of the flexibility due to overly short gap leads to competition between the polymerase and CoPrimer capture sequence for space. Additionally, upon DNA polymerase binding, the polymerase extension is partially obstructed by the steric inhibition by the capture sequence. However, it appears that this impediment of CoPrimer binding and extension initiation is aiding in suppression of unwanted annealing and extension (mutant-specific CoPrimer annealing to and amplifying the wild-type template), which results in enhanced CoPrimer specificity.

Ultimate C CoPrimer designs and Penultimate B CoPrimer designs (Table 1) possess gap, priming, and capture lengths that allows them to perform efficiently and robustly in multiplex assays (Fig 4). Both L858R assay and V769-D770insASV assay (a 9 bp insertion) exhibited excellent linearity down to MAF of 0.001 %. However, the G719S assay only exhibited good linearity down to MAF of 0.01 %. The less-than excellent performance of the G719S assay is not unexpected. The L858R mutation is a T>G substitution, resulting in G:A mismatch in the assay. The G719S mutation is a G>A substitution, resulting in G:T mismatch in the assay (in our G719S assay, the differentiating CoPrimer is the reverse one). The G:A mismatches near the 3’-end of the PCR primer are known to reduce polymerase activity by as much as 10 000-fold whereas G:T and C:A mismatches are among the least inhibitory mismatch types [50]. Additionally, the region surrounding the variant is extremely GC rich. The ultimate and penultimate p.G719S differentiating CoPrimer designs had priming fragments of a very high GC content (70 % and 72 %, respectively). It prevented us from decreasing the priming fragment Tm to a desired value without losing required amplification efficiency, and the high GC content may have contributed to undesirably high priming fragment stability of the final design.

Testing of ultimate and penultimate L858R, G719S, V769-D770insASV and *β-actin* quadruplex assays on cancer-cell derived *EGFR* reference materials confirmed reproducible and linear detection of the lowest MAF available (0.1 %) for all three mutations. Despite some differences observed during testing of ultimate and penultimate designs on synthetic materials, the results obtained on reference materials were almost identical, with only marginally better performance of the ultimate CoPrimer designs. From a practical standpoint, allele-specific CoPrimer designs with the variant induced mismatch at the penultimate base of the priming fragment appear to be more forgiving regarding the exact Tm of the priming segment. As such, penultimate CoPrimers would be easier to design and should have greater rates of success than the ultimate designs.

Allele-specific PCR CoPrimer assays, described in this publication, exhibit specificity and sensitivity that is typically required for rare allele analysis in liquid biopsy samples. Thanks to the elucidation of optimal CoPrimer design characteristics, custom assays can be generated easily and in a very short time. As CoPrimers are amenable to multiplexing, status of several mutations can be monitored simultaneously in one well. A quantification control assay, such as *β-actin, RNaseP* or *GAPDH* can be included in the multiplex assay as well. Unique CoPrimer structure guarantees very low rate of false positives originating from primer-dimer formation, while their specificity minimizes non-specific amplification due to mutant-specific CoPrimer mispriming and amplifying the abundant wild-type DNA.

